# Python-Microscope: A new open source Python library for the control of microscopes

**DOI:** 10.1101/2021.01.18.427171

**Authors:** David Miguel Susano Pinto, Mick A Phillips, Nicholas Hall, Julio Mateos–Langerak, Danail Stoychev, Tiago Susano Pinto, Martin J Booth, Ilan Davis, Ian M Dobbie

## Abstract

Custom built microscopes often require control of multiple hardware devices and precise hardware coordination. It is also desirable to have a solution that is scalable to complex systems and translatable between components from different manufacturers. Here we report Python-Microscope, a free and open source Python library for high performance control of arbitrarily complex and scalable custom microscope systems. Python-Microscope offers simple to use Python-based tools, abstracting differences between physical devices by providing a defined interface for different device types. Concrete implementations are provided for a range of specific hardware and a framework exists for further expansion. Python-Microscope supports the distribution of devices over multiple computers while maintaining synchronisation via highly precise hardware triggers. We discuss the architecture choices of Python-Microscope that overcome the performance problems often raised against Python and demonstrate the different use cases that drove its design: its integration in user facing projects, namely in the Microscope-Cockpit project; in controlling complex microscopes at high speed while using the Python programming language; and as a microscope simulation tool for software development.

## Introduction

The most advanced methods and applications in modern microscopy methods often require the assembly of custom built specialised microscope systems. Such one-off systems may range from an off-the-shelf microscope stand with a range of specialised devices attached, to an advanced engineering project built from custom optical components. There are many unique examples in the literature of such systems. When building most such systems, software development to control a specific combination of hardware devices forms a large portion of the effort. Other than developing software from scratch, there are currently three popular routes for controlling custom microscopes: individual device vendor software unique to each manufacturer; microscope specialised control software with the most popular being Micro-Manager (μManager), a general Java based framework with a GUI that has the lowest barrier to use; and custom control software often in LabVIEW or Matlab, both proprietary software. LabVIEW offers a visual programming environment that is commonly used for building instruments in the physical sciences, whereas Matlab is a programming environment with a focus on numeric computing.

The solution requiring the least specialist knowledge is to use the device vendor software for the individual microscope components and run them in parallel. For example, one would open the light source, the camera, and the stage programs separately, and place their individual windows side by side on the monitor to easily access all the controls. This solution is very convenient for simpler microscopes but lacks flexibility and scalability. Because the individual devices are being controlled on separate programs that do not communicate with each other, control of the microscope is effectively manual. This limits microscope usage to experiments that do not require synchronisation between devices. It also becomes extremely unwieldy; as more devices are used each inevitably requires its own separate program and window. Finally, with very rare exceptions, although these programs are often distributed at no cost they are not open source software.

μManager is a general free and open source program for control of microscopes (*Edelstein et al*., 2010). It is written in the Java programming language as a plugin for ImageJ (*Schneider et al*., 2012) and has a C++ core for hardware control that supports a wide range of hardware. More recently, Pycro-Manager has also been developed to provide control of μManager from Python and provide different hooks into a data acquisition pipeline for integration with image analysis (*Pinkard et al*., 2021). Having started in 2005, μManager continues under active development with a very wide and engaged user base, especially in the life sciences microscopy field. Its enduring development and widespread use is a testament to its quality and usefulness. But as discussed on *Chhetri et al*. (2020), μManager has limitations in setups with multiple cameras, its integration with hardware triggers, and more complex imaging modalities; can be difficult to create scripts, especially in complex configurations; and the combination of C++ and Java with an old tool chain required to build the software adds to the complexity of developing and debugging device adaptors, and of extending functionality. The migration of many scientists towards the Python programming language increases these issues. However, the widespread success of μManager means it has extensive hardware support and is often a good choice.

LabVIEW is a visual programming language and development environment that is widely used in physical sciences for prototyping instruments. It has very wide hardware support with many manufacturers providing LabVIEW modules (known as virtual instruments or VIs) to control their devices. Moreover, it allows very rapid development of graphic user interfaces with no previous programming knowledge. LabVIEW is proprietary and requires a commercial license for every computer it is used on, likely with associated cost. As each LabVIEW module is written by the equipment manufacturer there is little consistency in how to interface different hardware. The visual nature of the programming environment makes simple projects easy but systems with a large number of hardware components or complicated control architecture can become hard to understand, reproduce, and maintain. Although this complication can be reduced with good programming practices, it is not uncommon to outsource such work to a commercial company (*Chhetri et al*., 2020) because good code writing in LabVIEW is significantly more challenging than in popular general purpose languages such as Python. Additionally, the LabVIEW work flow does not integrate well into modern distributed source control infrastructure such as mercurial or git, a necessity for modern open source development.

Matlab is a numerical focused programming environment that scientists are often familiar with for data processing. It has frequently been used for microscopy, leveraging a number of available Matlab sub packages to provide GUI’s and easy access to complex data processing steps. The use of Matlab for microscope control is common in the field but the actual code is rarely shared and often custom to a single microscope setup and associated to image reconstruction (*Chmyrov et al*., 2013, *Ta et al*., 2015). Exceptions are ScanImage for the control of laser scanning microscopes (*Pologruto et al*., 2003), and Matlab Instrument Control (MIC) for the control of individual microscope components (*Pallikkuth et al*., 2018). Matlab provides a textual programming language simplifying code sharing and version control, however, Matlab is proprietary closed source software and the general requirement of many extensions significantly adds to the cost of implementing many systems.

There is currently an increasing number of software options for microscope control in Python, many of which are in the form of custom scripts specific to a microscope (*Alvelid and Testa*, 2019, *York et al*., 2013) but some provide a fully integrated microscope control environment, namely PYME (https://www.python-microscopy.org/) for SMLM and ACQ4 (*Campagnola et al*., 2014) for electrophysiology. While this code is freely available and can be modified, their design around a specific setup, technique, or environment reduces its potential for code reuse in other projects.

Several microscope vendors, such as Abberior Instruments and Zeiss, provide Python interfaces to enable instrument control from Python. These are all very useful additions to proprietary systems, however, they have a fundamental drawback that each manufacturer produces their own abstractions meaning code from one system is not compatible with another. Although these interfaces leverage the substantial Python infrastructure they are not generalisable and hence fail to enhance portability or reproducibility.

The fact that these companies are providing Python interfaces to their instruments indicates the general interest of the community in Python as a programming language to extend hardware capabilities. This demonstrates the potential benefit of an entirely Python based interface to a wide range of hardware.

In order to provide an alternative route that mitigates against these limitations, we have developed Python-Microscope, a Python library that provides an abstracted high level interface to control microscope devices. We have developed the software in Python, a programming language with which many researchers are already familiar, reducing the barriers to entry. Additionally, Python is a scripting language distributed as human readable plain text files, is cross platform, and does not require complex tool chains with compilation and linking steps, making it easier to share and reuse. Python also has a wide range of freely distributed packages for image analysis, which reduces the cost and complexity of integrating analysis with the control software.

Beyond the defined interface for each device type, Python-Microscope is designed to support arbitrarily complex microscopes with multiple devices spread across any number of computers. Being written in Python, a simple yet very powerful language, it enables complex and novel experimental approaches to be rapidly scripted and easily implemented.

## Library features

The design of Python-Microscope was based around a series of use cases derived from the experience of microscope hardware and software developers.

- Code independent from specific device models and vendors;
- Python programs as microscope experiments;
- Distribution of devices over the network and horizontal scaling;
- High performance hardware trigger and software trigger integration;
- Support both software and hardware developers during the development phase.

Here we describe these use cases and how Python-Microscope addresses them with the implementation details being described later.

### Write once, run with any supported device

A major design consideration is the ability to reuse code upon modification or reimplementation of a given microscope. This happens frequently: when replacing a specific device with a different model, either because the device failed or to get new features such as faster acquisition rates; when building a copy of the microscope on another site which has access to different devices; or as part of a larger improvement to an existing system, such as addition of a second light path with another camera. The ability to reuse the code after such changes to the system is critical because it accelerates microscope development by eliminating or reducing the software development work, which is incidental to the main purpose of the microscope.

At its core, Python-Microscope is a Python package that provides a defined interface for the different device types that may be used on a microscope. This means that code written for a specific device type, such a camera or light source, will work for all devices of that type. For example, to control the power output of a light source manufacturer-supplied control software for different devices not only uses different commands and names — such as power, intensity, or flux — but also different values — such as percentage, 0–1 range, or intensity in milliwatts. In Python-Microscope, the interface for light sources specifies the “power” attribute with a value in the 0–1 range for all light sources no matter what the underlying device implements.

A simple example of Python-Microscope being used in this way is the BeamDelta program (*Hall et al*., 2019) for alignment of optical systems that uses Python-Microscope to interface to any supported camera. Similarly, Microscope-AOtools (*Hall et al*., 2020) implements different methods of adaptive optics and exploits Python-Microscope to handle the control and specifics of different cameras and deformable mirrors to provide virtual devices. Python-Microscope is also used to interface hardware in Cockpit (*Phillips et al*., 2021) (see Supplemental Figure 1) and PYME (https://www.python-microscopy.org/), programs that provide a graphical user interface for complete microscopes and support a wide range of imaging modalities and a number of devices.

### Python programs as microscope experiments

The ability to write programs that are effectively microscopy experiments was an important goal. There are three reasons for this: First, to provide as much flexibility as possible, since graphical user interfaces are inherently limited by specific predesigned experiment types; Second, to enable feedback of arbitrary image analysis into any step of the experiment control; Third, to simplify the sharing of even extremely complex experiments. The ability to write experiments as programs provides all these features.

As Python-Microscope is a software library for controlling individual devices, it automatically supports the ability to create programs that are experiments. We have developed Python-Microscope so that such programs are easy to read and write, and even complex experiments can be encoded in easy to understand code (see examples in the documentation at https://github.com/python-microscope/). Python also provides an interactive environment, making development and debugging of systems as well as prototyping of experiments easily accessible.

### Arbitrarily complex mixture and number of microscope devices

Many novel microscopes require a large number of devices, sometimes reaching a point where it is not feasible to keep all devices connected to the same computer. This can happen either because many devices require too many resources, such as multiple cameras overloading the data bandwidth of the system, or because different devices have incompatible system requirements, such as requiring different operating systems.

For such situations, Python-Microscope was designed to support remote procedural calls and includes a *device-server* program which turns each individual device into a resource accessible over the network. A program can then access remote resources as if they were local Python objects. This design effectively makes the microscope a distributed system with a client-server model where each device is a server and the control program, typically a graphical user interface, is the client (Figure 1). This happens even if all devices are connected on the same computer since the client can connect to the server either locally or remotely.

**Figure 1:**
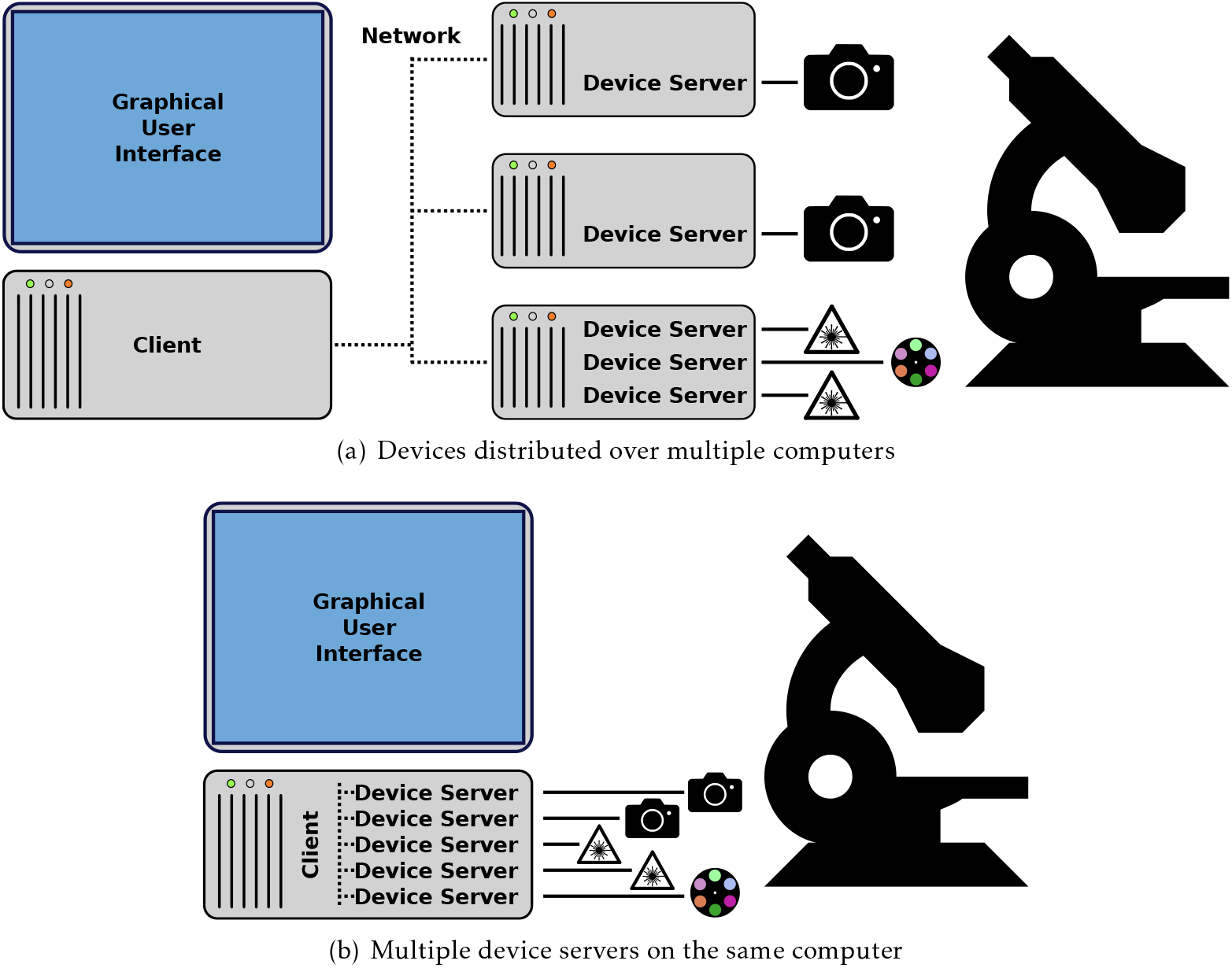
The device server: A client-server model enables the distribution of devices over multiple computers (a) and does not exclude the simpler case of having all devices controlled from the same computer (b).

CryoSIM (*Phillips et al*., 2020) and Aurox AO (*Hussain et al*., 2020) are two example setups that use the Python-Microscope *device-server* for hardware control in different ways.

CryoSIM incorporates four lasers and two filter wheels on one computer, a spatial light modulator on a second computer, two cameras on a third computer, and a fourth computer for overall system control, utilising the device-server to distribute many devices over several computers. The Aurox AO setup has all the device servers on a single computer, however the device-server model allows the integration of the Aurox Clarity module and a camera into a single composite device. Allowing the image processing to produce a widefield or confocal image from the raw image data to be integrated with image capture.

### Hardware and software trigger integration

Microscopy often requires strict timing. This is true even when imaging at low speeds because synchronisation is needed between multiple devices in order to have consistent exposure time at the expected moment. For example, exposure of a sample to the light source should be minimised to reduce photobleaching and photodamage so exposure should only happen while the camera sensor is active. Similarly, it is important to ensure that the time between start and end of light exposure is constant for reliable comparison between experiments. Controlling these devices with commands from a program, especially one running on a modern multi-tasking operating system, adds delay (latency) and reduces performance. In addition to the delay caused by the communication itself, there is unpredictable delay (jitter) introduced by the operating system context switching and asynchronous communication protocols, such as networks and USB. In total, these unpredictable timing issues can introduce changes of tens of milliseconds, higher than some experimental camera exposure times, making software control impossible for such demanding experiments.

Many devices can be configured to act on receipt of an electrical signal, often a TTL signal voltage, referred to as a hardware trigger. This hardware-based control is deterministic as it does not depend on any software. For instance a laser only emits light while receiving a high TTL input signal, a camera starts an image acquisition after receipt of a TTL rising edge signal, or a deformable mirror applies the next queued pattern. A device may also generate a trigger signal. For instance, cameras and light sources can often be configured to output TTL signals while exposing or emitting light respectively.

The microscope interface was designed with the concept of triggers that activate the individual devices and software triggers are handled as simply another trigger type. This approach provides an interface that supports software triggers but is easily upgraded to hardware triggers. The source of such hardware triggers can be other devices — typically a camera — or a dedicated triggering device. The recommended procedure is to prepare an experiment template that is then loaded on a dedicated timing device which triggers all other devices, as described in *Carlton et al*. (2010).

The existence of fast and cheap microprocessors and single board computers mean providing a dedicated hardware timing to sequence and synchronise a number of devices is relatively easy and extremely cost effective. We would recommend systems are designed around using an external device to provide hardware triggers to devices. This provides reliable timing and much more flexible sequencing than directly connecting outputs from one device to trigger inputs for another.

### Development tool

Finally, Python-Microscope is a useful tool during the system development phase. There are two sides of system development: the development of the actual optical setup, and the development of the software to control that optical setup.

To support development of optical systems, Python-Microscope includes a simple GUI program for the different device types (Figure 2). These are purposely kept simple as their aim is to test the basic functionality of the device without the added complexity that a full microscope control GUI brings.

**Figure 2:**
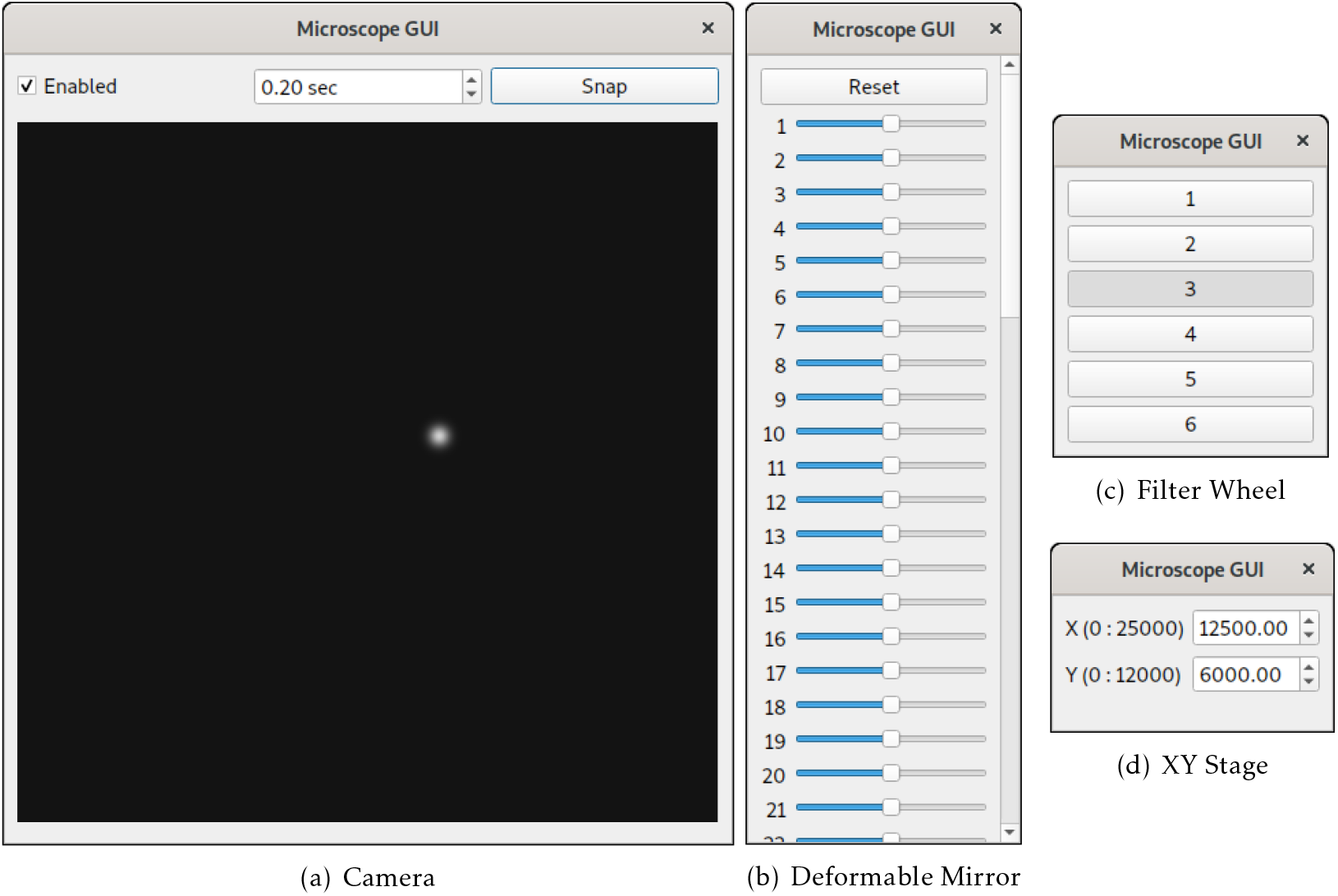
The *microscope-gui* windows provide a minimal GUI for the different device types to be used as development utilities. (a) GUI for a simulated camera that acquires single Gaussian shaped spots at random locations. (b) GUI for a deformable mirror provides a slider for each individual actuator. (c) GUI for a filter wheel displays a group of push buttons for the different filter wheel positions. (d) GUI for a simulated stage displays the different axis limits and positions.

To support development of user facing software, Python-Microscope provides simulated devices that enable software development without physical access to the hardware. For example, the simulated camera device can be configured to return images with specific features such as 2D Gaussian spots (Figure 2(a)) which were used during development of BeamDelta (*Hall et al*., 2019). Interaction between devices enables simulation of complex behaviour. For example, Python-Microscope provides a set of test devices to model sample navigation: a simulated camera uses the position of a simulated stage to “acquire” an image which is a subsection of a larger image. Focal position is simulated by blurring the image based on the position of the z axis of the simulated stage, and a “channel” is selected based on the position of the simulated filter wheel.

With hardware and software development happening concurrently, it is practical for one person to build the optical system and connect hardware while another is building the software. In doing so, total build time and costs can be reduced.

## Implementation

Documentation for the software and usage examples are available online at https://python-microscope.org. Here we include an overview of the design and implementation details as well as suggested workflows for hardware control using Python-Microscope.

### Abstract Base Classes

Python-Microscope interfaces for the different device types are implemented as Abstract Base Classes using Python’s abc standard library. These device ABCs are incomplete implementations with concrete methods that handle input validation and provide software fallbacks, and abstract methods for the individual device operations.

Currently, ABCs have been implemented for cameras, light sources, filter wheels, stages, and deformable mirrors, as well as integrated controllers that can control multiple pieces of hardware. All these ABCs inherit from a Device ABC, which defines core operations such as device settings and shutdown. The interface to support hardware triggers is implemented via an ABC mixin, named `TriggerTargetMixin`, which is mixed with the individual device type ABCs.

### Concrete Implementations

The implementation of a concrete class, i.e., support for a specific physical device, is device dependent but typically it either wraps a vendor-provided C library or a direct communication link to the hardware. In the case of a C library, these are dynamically loaded with Python’s ctypes standard library which removes the need for a compilation step or dependence on other tools. Direct communication is typically performed via RS-232 serial connections using Python’s pySerial package (https://github.com/pyserial/pyserial), frequently over USB to serial bridges in connected hardware.

Concrete implementations have been created for multiple devices. The full list is expanding with each new release and is part of the Python-Microscope documentation available online (https://python-microscope.org).

### Device Server

The *device-server* program creates a Pyro4 daemon (https://pyro4.readthedocs.io/) for each device. The Python-Microscope device instance is created and then registered with its Pyro daemon. There is only one Python-Microscope device instance per physical device. The Python multiprocessing standard library is used to place each Pyro daemon in its own python process, effectively sidestepping Python’s Global Interpreter Lock (GIL). The device server can be configured to serve a set of devices on the same process in order to simplify inter-device communication, typically in the case of composite virtual devices. Other instances that provide control of the hardware, such as individual stage axis and devices behind a controller, are registered with the Pyro daemon at initialisation. With Pyro’s autoproxy feature, these other instances are automatically replaced by a Pyro proxy on the client side.

While the use of device server is recommended, its implementation is decoupled from the device classes themselves and its use is entirely optional. This allows simple actions like debugging or testing partial systems to be performed directly via an interactive Python shell.

### GUI

The *microscope-gui* program makes use of the QtPy package (https://pypi.org/project/QtPy/) which abstracts the difference between PyQt5, PyQt4, PySide2, and PySide, all different Python wrappers to Qt, thus making the dependency a choice of the user. The dependency on QtPy itself is optional, making Python-Microscope easier to install if the GUI is not required.

For each device ABC there is a corresponding QWidget class. This widget is the central and only widget of the *microscope-gui* program window. The widget classes are public, so they can be incorporated in other programs. It should be noted that *microscope-gui* is not intended as a generalised user interface to control microscopes, we expect it to be used for low level hardware control under other software like Microscope-Cockpit (Supplemental Figure 1).

### Device specific features

An inherent limitation of defining an interface for different device types is the use of specialised device features which are specific to single device model or range of models. For example, some cameras have the option of applying denoising filters during acquisition, or provide control over amplifiers in the different stages of the camera readout. These features are exposed via a general “settings” mode, effectively a map of feature names to values. Python-Microscope provides such settings even when they clash with properties that are part of Python-Microscope’s interface. For example, Python-Microscope’s camera interface has a binning property but instances of `PVCamera` might also have the `BINNING_SER` and `BINNING_PAR` settings. Direct use of such settings, instead of the defined interface, ties the resulting code to the specific device, reducing potential for reuse hence we recommend against such practice if possible.

### Requirements

Python-Microscope has simple computational requirements. It requires Python 3.6 or later, and the Python packages NumPy (*Harris et al*., 2020), Pillow, Pyro4, hidapi, and pySerial. These are all free software and widely available for all operating systems, ensuring that Python-Microscope is truly cross platform. We have run Python-Microscope on different versions of Microsoft Windows, different GNU/Linux distributions, and macOS, on a multitude of computers from high-end workstation computers and rack servers, to touchscreen notebooks and single-board Raspberry Pi computers.

Despite the simple requirements that Python-Microscope itself has, individual devices often have their own more demanding requirements. Some devices require specific hardware connectors such as PCIe slots of a specific mode or size which can limit the motherboard or computer chassis used — for example, laptops and all-in-one computers very rarely contain any PCIe slots which are required for high speed and resolution cameras. Other devices may require a serial port or USB of a specific version. Similarly, use of devices at high speed will add further system requirements — e.g., acquiring many images at a fast rate will require faster data transfer and more computer memory. Finally, some of the device SDKs which Python-Microscope wraps are only available for one operating system, usually specific versions of Microsoft Windows. Note though that by using a network connection to the *device-server*, a controlling client program can be on another computer with a different operating system from the *device-server*.

### Workflow

Python-Microscope provides an API to microscope devices of different types. Their documentation is included as part of the class docstrings. While any program can easily instantiate the class of their specific hardware, our recommendation is to use the *device-server* program to create the devices and have programs control the devices via Pyro proxies. This removes the need for device specific code on the program, replacing it with configuration files that only need the device type and Pyro URI, both of which are strings that can be encoded on a plain text file. In the specific case of camera devices, the program needs to register the data client with a Pyro daemon and pass its URI to the camera device instance. This enables asynchronous transfer of images back to the controlling process.

The *microscope-gui* works as a simple example of the use of Microscope in a program for microscope control. The program is called with two arguments, the device type and its URI, a simpler form of configuration. Its CameraWidget class is also a minimal example of instantiating a queue object and serving it on its own dedicated Pyro daemon to be used as data client for the camera device.

## Discussion

Python-Microscope is a library to enable builders of custom microscope systems to fully exploit the power of the increasingly popular Python programming language. It interfaces with a growing range of hardware classes such as cameras, filter wheels, light sources, deformable mirrors, and stages with defined APIs, while still allowing access to device specific features. Python-Microscope provides simple access to complex devices for Python programs, and a defined protocol for accessing devices either locally or over network connections via the device-server. The device-server deals transparently with hardware restarts through reinitialisation and other issues, providing a reliable and robust interface to hardware in complex systems and environments.

Python is a free, modern, open source, portable programming language. It has wide support and a large developer community, it is relatively easy to learn, and has a wide ecosystem for scientific computing. However, Python is not without its drawbacks. In the context of hardware control, namely the coordination of multiple hardware devices, the most commonly mentioned issue is Python’s performance and lack of “true” multithreading due to a Global Interpreter Lock (GIL).

As mentioned in the introduction, μManager has recently introduced a Python interface, Pycro-Manager (*Pinkard et al*., 2021). This simplifies connections between μManager based hardware interfaces and Python based analysis and control. Although this reduces the effort in using Python for control and online analysis compared to other approaches, it does not provide direct access to the hardware via Python and keeps the existing μManager infrastructure, namely that knowledge of both C++ and Java is still required for development and debugging.

Python has the reputation for being slow when compared to C, C++, or even Java, apparently making it an odd choice for microscope control where multiple physical devices have to be controlled at high speed in an orchestrated manner. However, multi-threaded operating systems simply do not have reliable enough timing for the triggering of critical components, whichever programming language is used. For these applications, we recommend the use of hardware trigger modes that many devices have (or are available as options). This approach was one of the main design considerations for Python-Microscope, as it makes Python’s performance mostly irrelevant to the overall performance of the microscope system. Moreover, we recommend the use of Python-Microscope in a client-server model to access the different devices, each of them with a dedicated Python process, thus side stepping the GIL.

Even if hardware triggers and a client-server model are not used, our experience is that Python’s reputation as being slow is unjust. At least in the case of scientific computing, the core processing steps in Python are often implemented in C or Fortran. In Python-Microscope, most of the hardware controllers are simply wrappers to vendor provided libraries themselves written in C. In addition, calls to functions in those C libraries are done with Python’s ctypes which also releases the GIL.

Python has become the de facto standard in the field of data science. CellProfiler, Ilastik, NumPy, PyTorch, SciPy, and TensorFlow are well known examples of Python programs and libraries. Novel advances in the field of machine learning are increasingly happening first in the Python programming language. By using the Python language, Python-Microscope can harness all of these and bring new developments into the field of microscope control more quickly and easily, accelerating the advancement of the field to what we see as the next logic step (*Waithe et al*., 2020).

A further advantage of the approach provided by Python-Microscope is in increasing reproducibility in science. Scientists frequently write in their publications that LabVIEW or a specific commercial microscope acquisition software was used without any specific acquisition settings, code, or macros to assist with reproduction. This is especially critical in complex experimental setups where specifics of acquisition are particularly important. Python-Microscope has the potential to change this trend, allowing authors to freely publish simple code demonstrating exactly how their control and acquisition code operates. Additionally, the defined device interfaces allow such code to be ported to other specific hardware with minimal changes.

Python-Microscope is a free and open source project currently being used in several labs with an open development approach. Our aim is that the microscope development community will find it a useful tool and engage in this development to increase its general usefulness. With that aim in mind, we perform our development conversations and user support in the open as github issues and the project is an image.sc community partner. In particular, expanding the number of devices supported by Python-Microscope would be extremely beneficial. However, adding support for a device requires physical access to the device and the current list of supported devices echoes the devices we and our collaborators have access to. This is a chicken and egg problem. Python-Microscope needs broad device support to be widely adopted by the community but it needs contributions from the community to support those devices. We believe that Python-Microscope currently provides enough devices and infrastructure to support adoption by more developers. There are contribution guidelines within the “Get Involved” section of the documentation, available online at https://www.python-microscope.org/doc/get-involved. In an effort to minimise duplication and increase community cooperation we are keen to foster links and collaborations with other Python based, or Python compatible microscope control packages such as PYME and μManager. We will endeavour to make bindings, when possible, that will enable interoperability of Python-Microscope with other open community-based Python software controlling microscope hardware.

## Conclusion

We have developed Python-Microscope to sit at the foundation of a microscope control software stack. The foundation is formed by abstracting the differences between individual devices which dramatically reduces the differences in software between microscopes. By using Python, which has a large user base in the scientific community, we are able exploit the existing expertise in the field and leverage a vast range of scientific libraries for image analysis. The design and features simplify the reuse and sharing of software components between custom microscopes. We expect that it will ultimately accelerate and expand microscopy developments.

## Software Availability

Python-Microscope is distributed under the GNU General Public License version 3.0 or later. Source releases are available for download from the Python Package Index (https://pypi.org/project/microscope/). Development sources are available from GitHub (https://github.com/python-microscope/microscope). Documentation is included in the source releases as well as HTML on the project website (https://python-microscope.org/).

## Acknowledgements

We would like to thank members of Micron Oxford for helpful comments and suggestions during the development of Python-Microscope. We also thank Dominic Waithe from the University of Oxford, and Thomas Huser and his group from the University of Bielefeld for their help during testing and development of more devices. We are also thankful to David Baddeley, from the University of Auckland, for hints from his PYME package, suggestions, and discussion around hardware control from Python.

## Author Contributions

DMSP: conceptualisation, software, writing — original draft preparation, writing — review and editing. MAP: conceptualisation, software. NH: software. JML: software. DS: software. TSP: software. MJB: funding acquisition, supervision, writing — review and editing. ID: funding acquisition, supervision, writing — review and editing. IMD: conceptualisation, software, supervision, writing — original draft preparation, writing — review and editing.

