## Supplemental figures for "Python-Microscope: A new open source Python library for the control of microscopes"

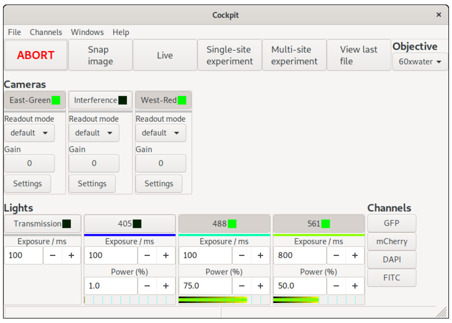


(a) Main Window

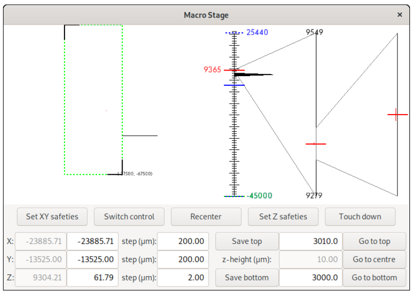


(b) Macro Stage

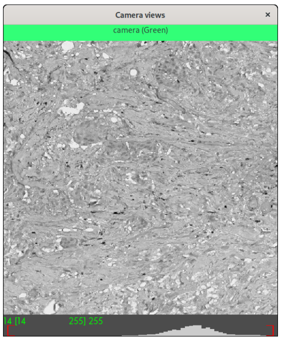


(c) Camera View

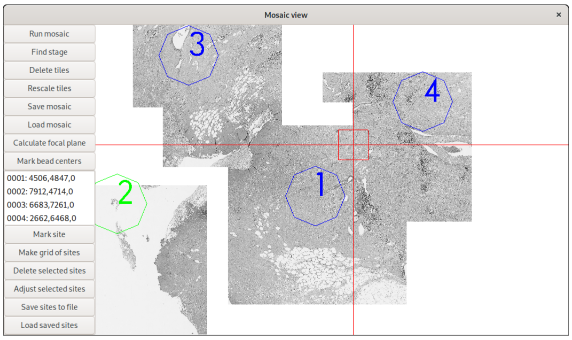


(d) Mosaic View

Supplemental Figure 1: An example of a complex GUI interface based on Python-Microscope. The Microscope-Cockpit GUI running a microscope with 3 light sources, 3 cameras, a XYZ motorised stage, and a Z piezo insert. Figure from *Phillips et al.* (2021). (a) The main Cockpit window provides quick access to a number of functions and shows the status of all connected devices, as well as the channels buttons to switch between pre-defined configurations. (b) The macro stage window provides an overview of the position of all stages, including nested Z stages. (c) The camera view shows the last image taken from each active camera and their histograms. (d) The mosaic view displays an overview of the sample. It saves images from tiles of mosaic scans onto the computers GPU. This gives the possibility to store hundreds or thousands of images as a single mosaic, which can be navigated from GPU memory in real time.

[1]Micron Advanced Bioimaging Unit, Department of Biochemistry, University of Oxford, South Parks Road, Oxford, OX1 3QU, United Kingdom
[2]IGH, Univ Montpellier, CNRS. 141 rue de la Cardonille, 34396 Montpellier, France
[3]Department of Engineering Science, University of Oxford, Parks Road, Oxford, OX1 3PJ, United Kingdom
